# Inferring metabolite states from spatial transcriptomes using multiple graph neural network

**DOI:** 10.1101/2024.06.12.598759

**Authors:** Jiaxu Luo, Daosheng Ai, Wenzhi Sun

## Abstract

Metabolism serves as the pivotal interface connecting genotype and phenotype in various contexts, such as cancer reprogramming and immune metabolic reprogramming. Compared to the transcriptome, the development of the single-cell metabolome faces significant challenges. While various methods exist for predicting metabolite levels from transcriptome, their efficacy remains limited. We developed an efficient and adaptable algorithm known as Multiple Graph-based Flux Estimation Analysis (MGFEA). MGFEA enables rapid inference from million-level single-cell transcriptome datasets and achieves accuracy comparable to that of scFEA. Additionally, MGFEA can detect metabolite biomarkers in different cancer bulk RNA-seq datasets. As an attempt to integrate multi-omics dataset, MGFEA can further improve the accuracy of these inferences by leveraging additional metabolome.

## Introduction

In the intricate realm of cellular biology, common cells diligently uphold metabolic homeostasis within their internal milieu, a crucial state that ensures the proper functioning of distinct functional proteins. Notably, the diverse nature of cells entails varied modes of sustaining metabolic equilibrium. For instance, neoplastic cells exhibit metabolic reprogramming, altering their metabolic profiles to adapt to their environment [1–5]. Similarly immune cells dynamically adjust to effectively adapt and respond to their microenvironment [6–8].

Over the last decade, there has been significant advancement in single-cell transcriptomics technology, leading to the accumulation of a substantial number of precise single-cell databases [9–12]. However, the advancement of single-cell metabolomics has lagged behind that of single-cell transcriptomics largely due to inherent technical bottlenecks [13,14]. Though progress has been slow, contemporary computational tools are now capable of characterizing metabolism through transcription. Enrichment-based methods have demonstrated significant impact in the field of functional genomics research, yet they are predominantly utilized for qualitative analysis [15–18]. Constraint-based models have shown significant promise by their ability to deduce the rate of metabolic reactions without the prerequisite detection of numerous kinetic parameters [18,19]. Consequently, novel computational models have emerged for predicting flux state at the single cell resolution. These include, but are not limited to, scFBA [20], scFEA [21], Compass [22], and METAFlux [23]. Despite their innovative nature, the efficiency of these models still falls short of optimal levels.

Building upon the foundations established by scFEA and a series of constraint-based models, we introduce a novel modeling framework for inferring metabolic flux based on both metabolic network guided gene interaction graph and spatial information graph (Fig. 1a). This framework aims to estimate the divergences in metabolic reactions among cells by utilizing the known gene-reaction relationships in the Genome Scale Metabolism model (GSMM) alongside single-cell transcriptomics dataset. The code implementation and relevant dataset are available on GitHub (https://github.com/Sunwenzhilab/MGFEA).

**Fig. 1:**
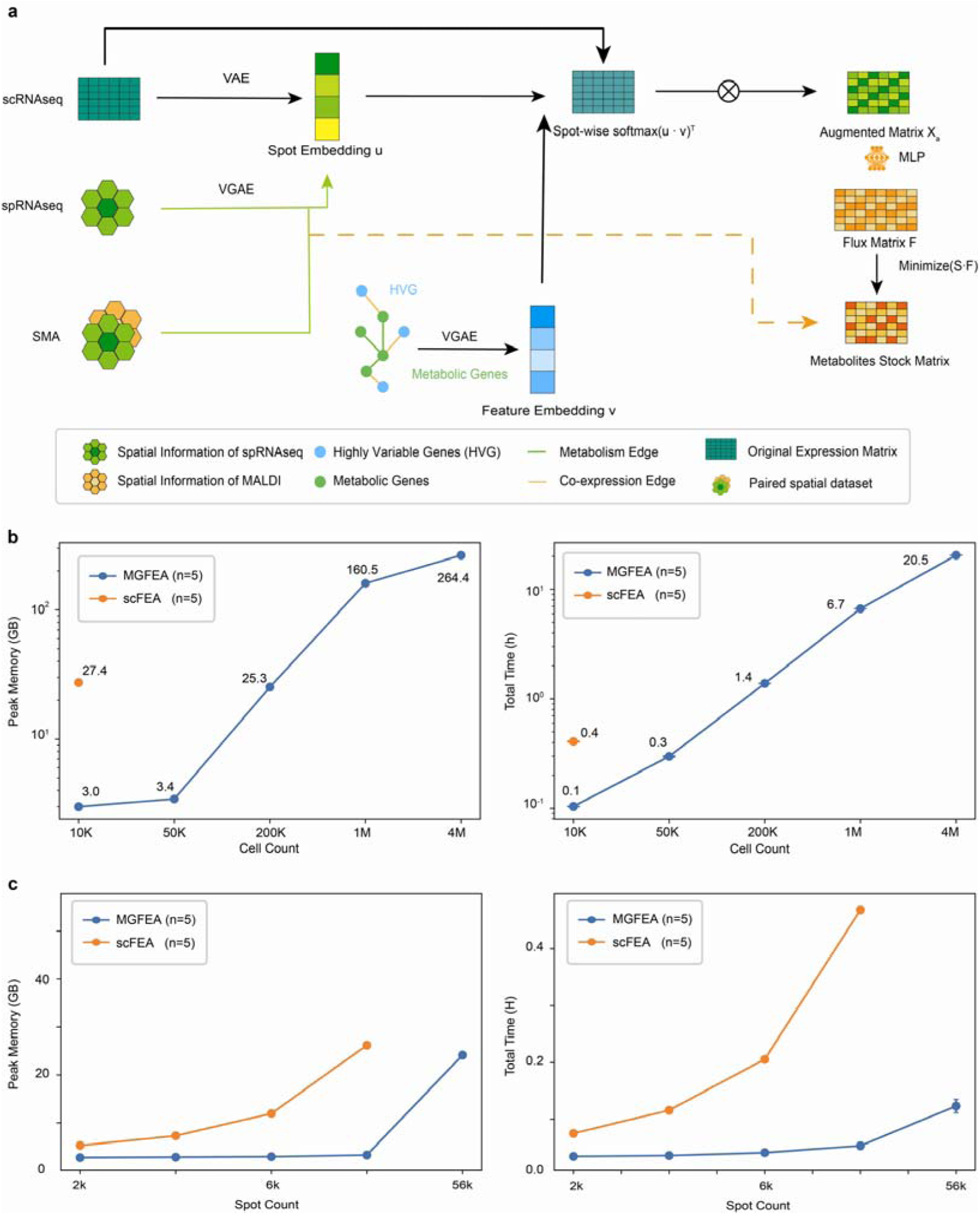
An algorithmic framework to estimate metabolic states based on spatial and single cell transcriptomic dataset, and computational performance measurement. a, MGFEA algorithmic framework. VAE: variational autoencoder VGAE: variational graph autoencoder scRNAseq: single cell transcriptome dataset spRNAseq: spatial transcriptome dataset MALDI: Marix-Assisted Laser Desorption Ionization SMA: spatial multimodal analysis dataset. A paired spRNAseq and MALDI dataset. b, left: Comparison of peak memory usage between scFEA and MGFEA for different sized datasets three repeats each point. right: Comparison of time cost between scFEA and MGFEA for different sized datasets three repeats each point. scFEA reported out of memory error in the 4 large datasets, so there is no data point of scFEA in the figure. c, left: Comparison of peak memory usage between scFEA and MGFEA for different formats of stereo-seq dataset. Right: Comparison of computation time between scFEA and MGFEA for different formats of stereo-seq dataset. scFEA reported out of memory error in the 4 large datasets, so there is no data point for scFEA in the figure.

## Result

### Overview of MGFEA framework

MGFEA pipeline combined the framework of representation learning and the metabolic flux constraint framework such as scFEA and adopts a suitable module for sparse matrix. MGFEA is a flexible framework which achieved the efficient and accurate metabolic inference for various types of datasets, such as single cell RNA seq dataset, spatial RNA seq dataset. MGFEA extracted cell embeddings from expression matrix and integrated gene co-expression information from expression matrix, spatial information from spatial RNAseq and expert knowledge from GSMM model into the gene embeddings. MGFEA used the dot product to combine the two types of embedding and used the dot product as weights to enhance the cell expression embedding. At the last layer, full connected layer transformed the enhanced cell expression embedding into the metabolic flux under the constraint of the metabolic flux loss. With the result of metabolic flux, MGFEA could infer all of the metabolites’ relative stock level in the single cell.

Compared with scFEA, MGFEA demonstrated three innovations. First, our framework exhibits remarkable flexibility, featuring multiple modules that can accept inputs in various data formats, including single-cell transcriptome datasets, spatial transcriptome datasets, and spatial multimodal analysis (SMA) datasets [24]. MGFEA employed anndata format and sparse matrix module to achieve compatibility for large datasets. Second, MGFEA utilized representation learning to extract and integrate gene interaction information from expression matrix, spatial information from spatial RNAseq and expert knowledge from GSMM model. By leveraging Variational Graph Autoencoder (VGAE) or Variational Autoencoder (VAE), our framework extracts cellular embeddings from the transcriptome dataset. Additionally, another VGAE is employed to extract inter-gene metabolic information derived from the knowledge-guided graph of GSMM, thereby generating gene embeddings. We then computed the dot product of these two distinct embedding types and applied a softmax transformation along the cell axis, to derive a weight matrix. This matrix was subsequently applied to enhance the original transcriptome matrix. Third, based on the premise which non-metabolic genes also affect metabolic flux, MGFEA used the same stoichiometry matrix from scFEA and other GSMM models to guided fully connected multiple layer perceptron (MLP) to transform the extracted cell embeddings into metabolic flux in the cell.

To promote the development of metabolic prediction methods based on metabolic graph, we also provided three types of metabolic graphs for each species: the first from scFEA, the second from flux-estimator [42], and the third from the GSMM model Recon3D and IMM1865 [26,27]. Although flux-estimator [42] only subgraphs for user access, we integrated the majority of these into a single comprehensive graph to ensure a fair comparison. Graphs serve as an important role in the constraint based methods. In this article, when referencing the small graph, we will abbreviate the names of two models. For the large graph, we will append an "-L" suffix to the different models. Similarly, IMM1865 will be referenced to as "IMM." Subsequently, the improved matrix serves as input, enabling the neural network to autonomously learn without fixed gene-reaction relationships. Through unsupervised metabolic flux inference, our framework derives metabolic flux, facilitating the characterization of metabolite imbalance levels across different cells analogous to metabolomics analysis. Furthermore, owing to the flexibility of flux estimation analysis method, we also provided an additional reference module to leverage the metabolic information present in the SMA dataset thereby further optimizing the algorithm’s performance. Based on the existing contributions of scFEA [21], flux-estimator [42], Recon3D [26] and IMM1865 [27], we also offered three types of graph in h5ad format and the reference module can also be applied on the other constraint based methods. The above two items represent our modest contribution to the field of constraint-based inferences methods about metabolism.

### MGFEA outperformed in memory usage and time cost

Based on the data preprocess pipeline which is suitable for the sparse matrix, MGFA showed high computational performance. Compared with scFEA, MGFEA demonstrated significant advantages in computational performance, particularly in its adaptability to the growing scale of single-cell datasets and its efficiency regarding memory usage and computational speed (Fig. 1b). In the same device, scFEA pipeline reported out of memory error in the large datasets, but MGFEA showed measurable performance. Considering the scalability of stereo-seq datasets [43], we selected five thresholds, resulting in the generation of five different datasets sizes, each with varying sizes and library depth of spots. According to stereo-seq datasets, MGFEA demonstrated better performance than scFEA in different resolution of stereo-seq datasets (Fig. 1c). According to the benchmark of the daily largest single cell RNAseq datasets, MGFEA exhibited improved performance in terms of computational resource utilization.

### Depmap dataset benchmark confirms framework efficiency and discovery of potential metabolites biomarker

Metabolic reprogramming is the classical features of tumor. Tumor metabolic reprogramming is the important format of tumor rearranging tumor microenvironment and resisting the chemotherapeutic agents [44]. We validated the performance of MGFEA in the public datasets which owns transcriptome and metabolome information of human cancer cell lines. To assess the effect of dataset size to model accuracy, we utilized a public dataset paired with metabolomics dataset from human cancer cell lines provided by the Cancer Dependency Map (DepMap) Project [36,37]. We selected all overlapped metabolites among the different metabolic graphs and the metabolomics dataset for comparable accuracy assessment across different metabolic graphs. Then we calculated the relative mean square error between the predictions and metabolomics result. We presented the results of the prediction from scFEA, scFEA-L, MGFEA, MGFEA-L and MGFEA-IMM. As the dataset size increased, the performance of two flux estimation analysis algorithms improved. The usage of flux estimation analysis algorithm on larger datasets showed higher inference accuracy and the performance of predicted results from MGFEA consistently outperformed that of scFEA (Fig. 2a).

**Fig. 2:**
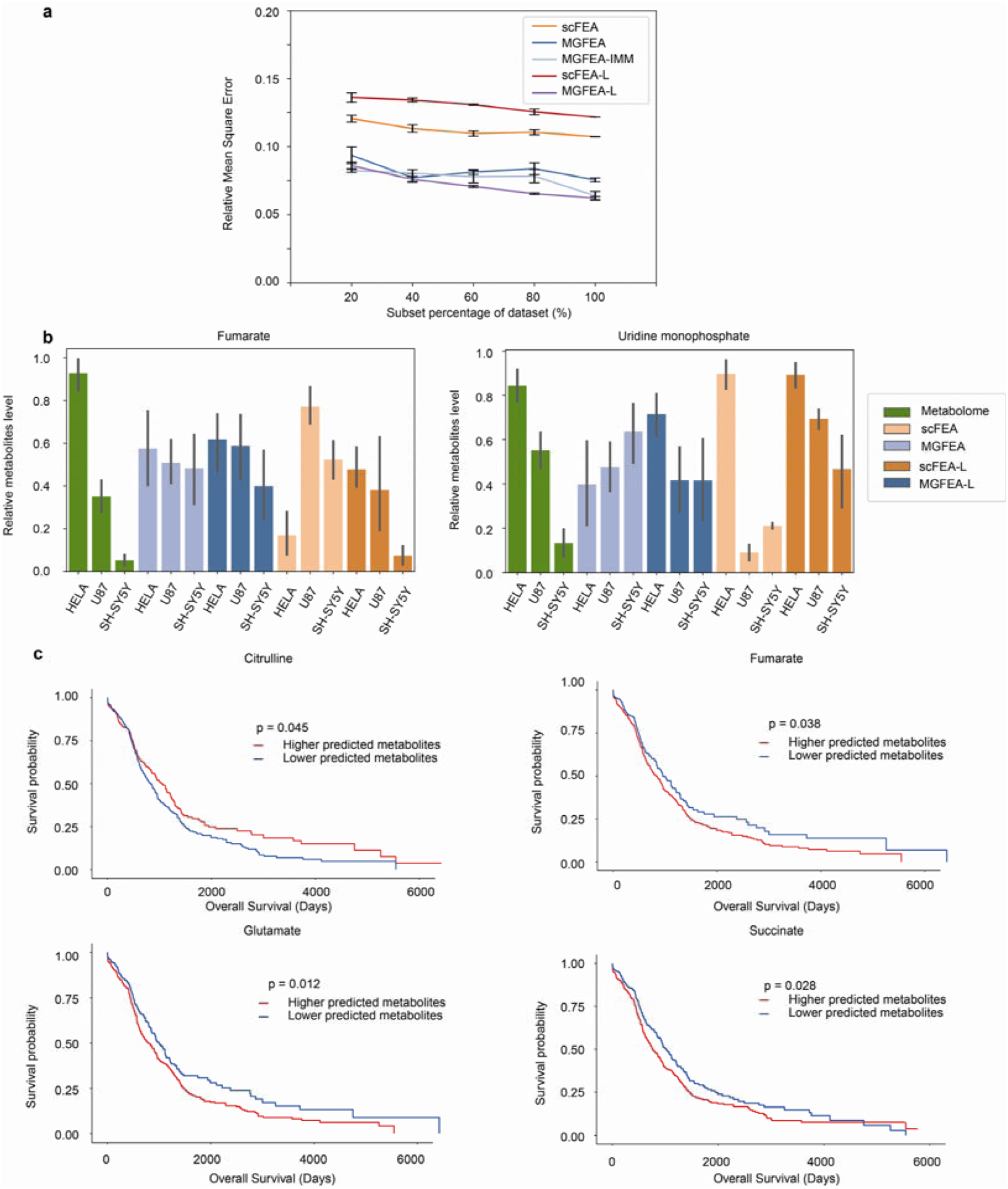
Comparison of prediction results from scFEA and MGFEA in DepMap dataset and predicted biomarker in TCGA glioma dataset. a, The relative mean square error comparison of metabolic predictions from scFEA, MGFEA, scFEA-L, MGFEA-L and MGFEA-IMM on DepMap datasets. b, The prediction result from metabolomics, scFEA, scFEA-L, MGFEA and MGFEA-L at Fumarate and UMP. c, The prognosis of TCGA glioma patients is correlated with several predicted metabolites: citrulline, fumarate, succinate and glutamate. Red lines mean the statistics from patients with higher predicted metabolites. Blue lines mean the statistics from patients with lower predicted metabolites.

**Fig. S1:**
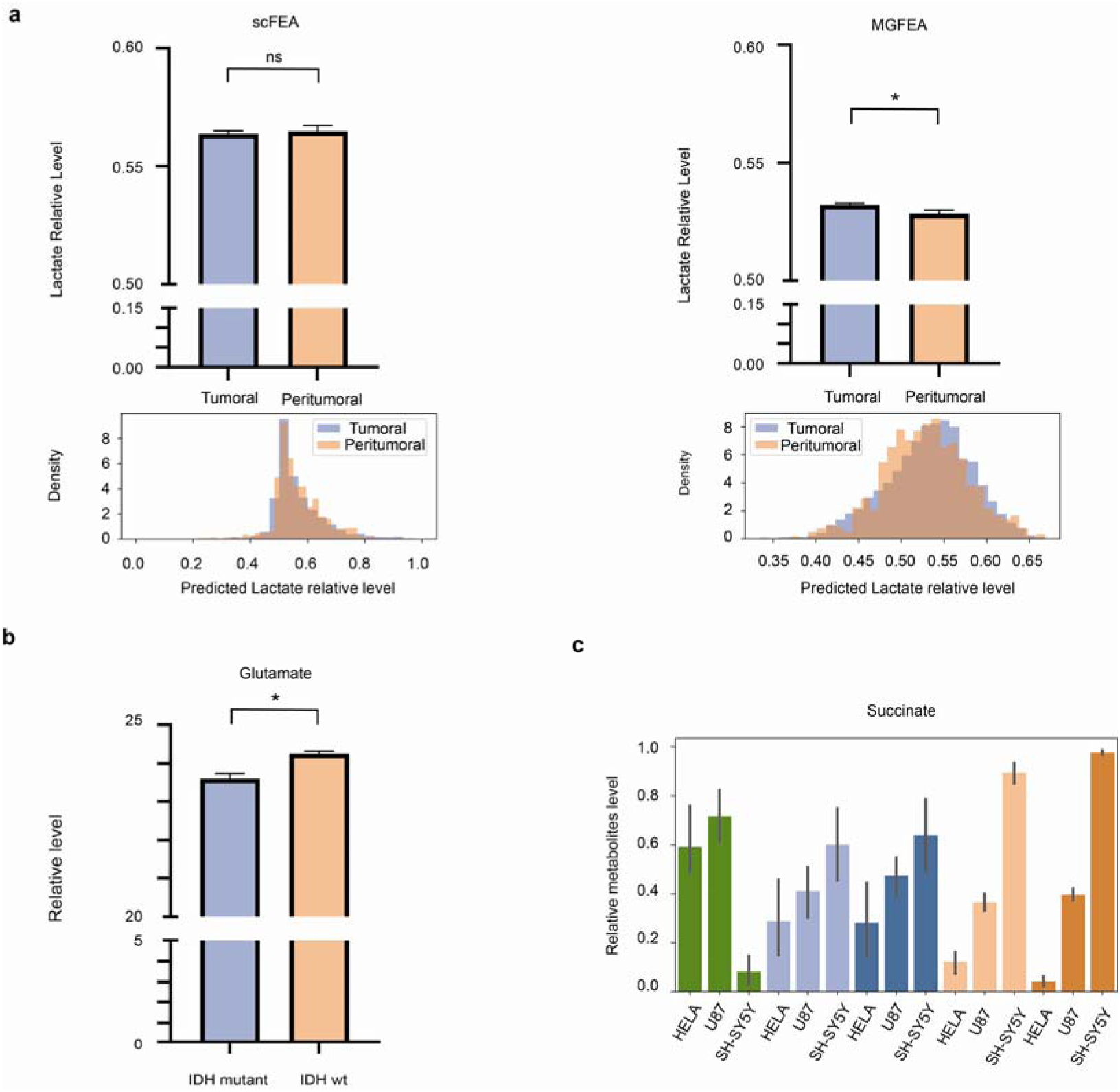
Comparison of MGFEA and scFEA in terms of cancer metabolites imbalance level. a, Predicted relative lactate levels of lactate in different glioma regions by MGFEA and scFEA. b, Metabolomics result about glutamate in IDH wild type and IDH mutant type glioma samples from Wang et al. c, Inhouse datasets metabolomics results and prediction results from scFEA and MGFEA about succinate content in Hela, U87 and SHSY5Y cell lines.

As a supplemental experiments, we cultured three types of cancer cell lines: Hela, U87 and SH-SY5Y for our in house paired transcriptome and metabolome dataset. Based on the targeted metabolomics detection, we confirmed that for the ten detected metabolites, the predictions from MGFEA-L and scFEA-L matched with the true concentration distribution in fumarate and Uridine Monophosphate across three cancer cell lines targeted metabolomics detection (Fig. 2b). But both MGFEA and scFEA show incorrect trends in other two significant differential metabolites: beta-alanine and deoxyadenosine. Two models reported the same predictions but not matched with the validation completely on succinate (Fig. S1c), we will discuss the phenomenon in the discussion section.

MGFEA could be used in the discovery of tumor potential metabolic biomarkers. In an application utilizing single-cell transcriptome dataset from gliomas sampled across different regions [45], we observed a significant difference in lactate levels between cells in the tumor core and those in the peripheral regions as predicted by our model (Fig. S1a). This finding is similar with the functional magnetic resonance imaging results [46]. Leveraging the abundant transcriptome datasets from TCGA [20], we tried to find the potential biomarker with the prediction from MGFEA. MGFEA classified of patients as two groups based on the median relative predicted metabolites level from MGFEA. For example, MGFEA identified four metabolites as the potential biomarkers that could distinguish patients with poor survival outcomes (Fig 2c). Citrulline is predicted as a biomarker associated with patients’ better prognosis (Fig 2c). Citrulline has been found as the products of nitric oxide(NO) synthase, which catalyzed the reaction which produce NO. Similar research supported our predictions that oral administration of L-arginine or hydroxyurea significantly increased brain tumor barrier permeability when compared with the nontreated control rat [47]. Thereby patients whose tumor owns higher citrulline level could be more sensitive to the chemotherapeutic agents. MGFEA predicted that fumarate is associated with patients’ bad prognosis. Fumarate has been validated that correlated with the inhibitory function of CD8 positive T cell [48]. Similar measured results about Glutamate can be found in a published metabolomics dataset [49] (Fig. S1b). Glutamate is found in higher concentrations in IDH wild-type gliomas but is lower in IDH-mutant gliomas. Our predictions are consistent with metabolomic findings, as IDH-mutant patients, who generally have better prognosis, show lower glutamate levels. Research has indicated that glioma release glutamate to improve their growth by utilizing its neurotoxicity [50,51]. MGFEA also predicted succinate correlated with the patients’ bad prognosis. Glioblastoma cells improved the succinate level to accommodate the hypoxic environment which is similar with our predictions [52]. MGFEA showed comparable accuracy with scFEA in DepMap datasets and could be used for the identification of potential metabolites’ biomarkers in TCGA and more human datasets.

### MGFEA integrated non metabolic genes and metabolomics into the whole framework

With the spatial multimodal analysis (SMA) paired spatial RNAseq and Matrix-assisted laser desorption/ionization (MALDI) datasets, we could validate the performance of MGFEA and validate the true metabolic information integration module of MGFEA. For the SMA dataset, we selected one of the slices to compare the prediction results of different models against MALDI results (Fig. 3a-d). The finding indicated that MGFEA captured spatial similarity of different spots and showed significant better performance when using the raw matrix as input. In both models of MGFEA, imputation did not yield significant improvements in performance (Fig. 3a, c). Consistent with scFEA reports, MAGIC [31] improved its performance (Fig. 3b, d). STAGATE [32] which integrates spatial information demonstrated no improvement in scFEA and small improvement in scFEA-L. We compared three methods which select variable genes for MGFEA metabolic gene interaction graph construction: common highly variable genes (HVG) defined by normalized dispersion, spatial autocorrelation metrics Moran’s I (SRR) and spatial differential expressed genes (SDE). By leveraging HVG, MGFEA demonstrated better performance than the other methods without imputation. Although HVG methods did not demonstrate improvement compared to the other two methods, they also did not result in decrement in performance. Therefore, we used HVG as the default parameter of MGFEA. With respect to the relative mean square error, MGFEA outperformed scFEA with different imputation preprocess (Fig. 3b, d).

**Fig. 3:**
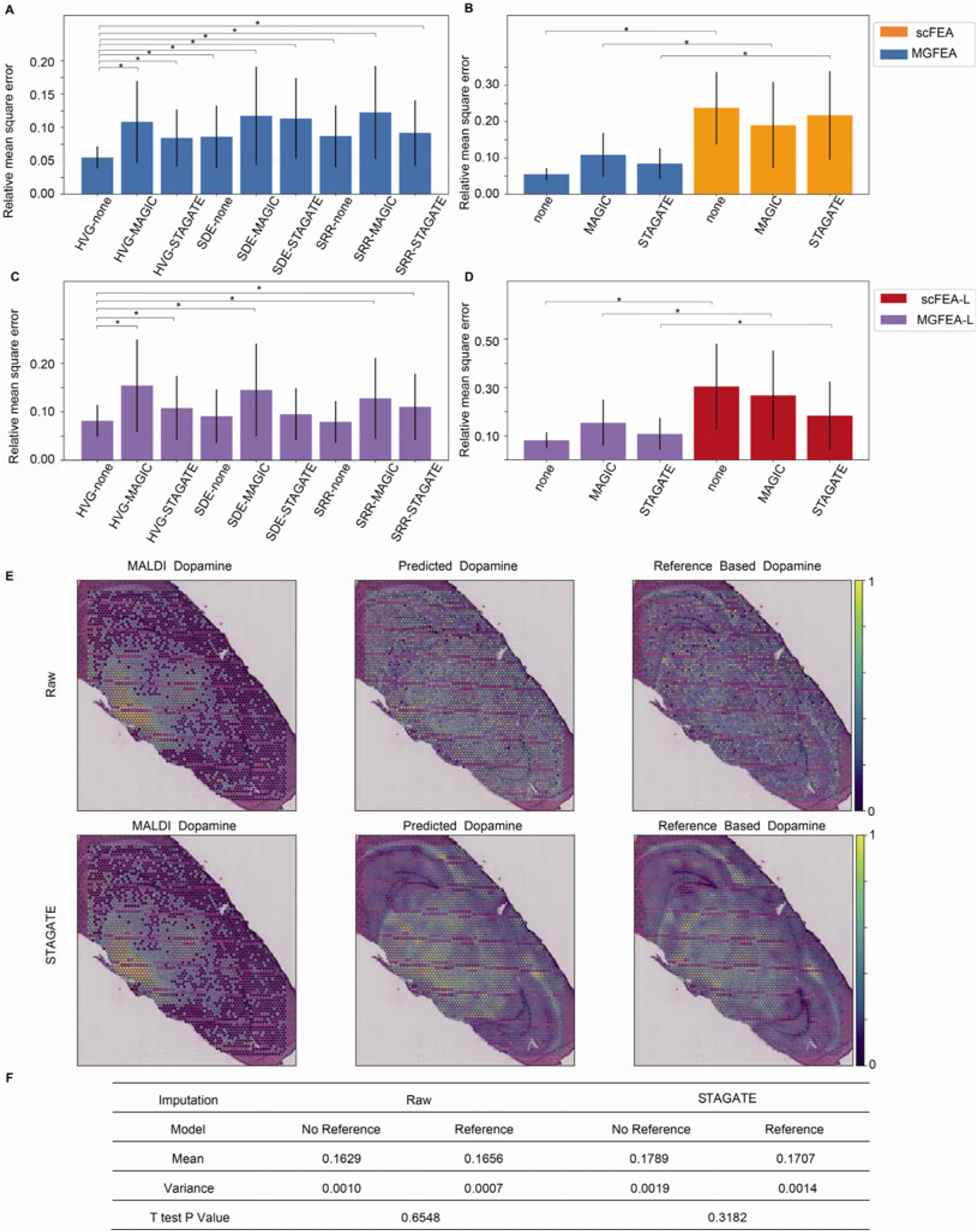
Comparison of prediction results from scFEA and MGFEA in SMA dataset and the reference module which integrates metabolomics and gene expression. a, c Relative mean square error between the prediction by MGFEA and MGFEA-L with different variable genes selection methods, highly variable genes, spatial correlated genes and spatial differential genes. b, d Relative mean square error between the prediction by scFEA, MGFEA, scFEA-L, MGFEA-L in SMA dataset with the three different imputation methods, no imputation, MAGIC and STAGATE. e, Comparison of dopamine prediction basic MGFEA and referenced MGFEA with raw matrix and imputated matrix as input. f, Statistics of reference module contribution in MGFEA relative mean square error.

Given the paired nature of the SMA dataset, leveraging the metabolomics results as a reference for MGFEA predictions can improve performance (Fig. 3e). We observed improvements in accuracy and reductions in variance when using the MALDI reference, although statistical significance was not achieved (Fig. 3f).

### Unpaired brain transcriptome and metabolome datasets confirmed efficiency of MGFEA

Leveraging the unpaired metabolome and spatial transcriptome datasets, MGFEA showed efficient performance on differential metabolites identification. Based on the metabolomics dataset of different brain regions from Shao et al [53] and visium sagittal spatial RNAseq dataset, we present the results of the metabolome, the prediction from scFEA, MGFEA, MGFEA-IMM, scFEA-L and MGFEA-L (Fig. 4a). Compared to scFEA, MGFEA showed lower relative mean square error across all metabolic graphs. We provided several examples of prediction results (Fig. 4b). MGFEA exhibited comparable prediction accuracy to scFEA in two types of metabolic graphs (Fig. 4c, d). Along with the significant differential metabolites identified through metabolomics, MGFEA demonstrated greater accuracy in classifying relative differences across various brain region (Fig. 4c). When considering the top-ranked predicted differential metabolites from each model, MGFEA exhibited outperformed scFEA (Fig. 4d).

**Fig. 4:**
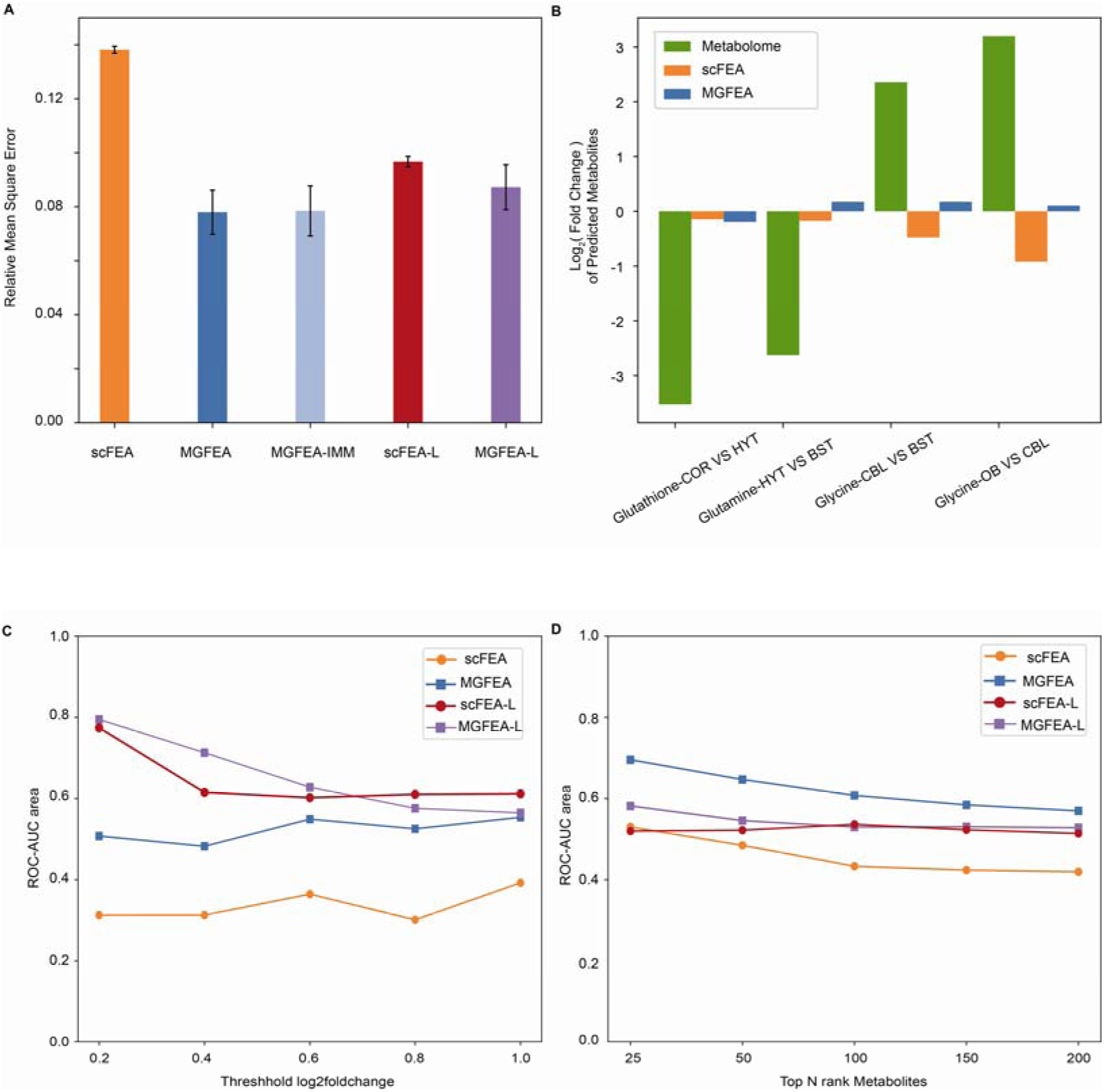
Comparison of prediction results from scFEA and MGFEA in unpaired visium sagittal dataset and Shao metabolomics dataset. a, Results of relative mean square error between Shao datasets and the prediction of scFEA, MGFEA, scFEA-L, MGFEA-L and MGFEA-IMM for the relative metabolite level of mouse brain. b, Detection of log fold change of residual metabolites from different brain region pairs using metabolome, scFEA and MGFEA. COR: cortex, HYT: hypothalamus, BST: brain stem, CBL: cerebellum, OB: olfactory bulb. c, The ROC-AUC area of models in relative difference classification within significant differential metabolites from metabolome. d, The ROC-AUC area of models in relative difference classification within top differential predicted metabolites from models.

## Discussion

MGFEA is designed for fast inference on large datasets and is particularly adept at inferring metabolic states in tumor samples, leveraging the rich transcriptomic public data resource. Our pipeline improved computational performance significantly and could be applied in the analysis of large datasets even million level datasets [54,55] and the future application on the insilico perturbation gene functional analysis which needs thousands repeats.

Tumor heterogeneity is associated with the bad prognosis of patients [56]. In contrast to the metabolome, transcriptome enabled researchers to acquire single cell transcript information at an affordable price [57]. Thereby we used MGFEA in tumor samples for the discovery of biomarkers. To validate the efficiency of models, we prepared the in-housed dataset of three cancer cell lines. In the in-housed dataset, both MGFEA and scFEA predicted the correct distribution of fumarate and uridine monophosphate. Interestingly, scFEA and MGFEA reported the same prediction about succinate based on the transcriptome, but different from the targeted metabolomics detections. There are several possible reasons about the phenomena, for example, the correlation between metabolites and transcript is weak [24]. Enzyme catalyzed metabolites transformation, enzyme is translated from transcript, but the correlation between protein and transcript is even weak [58]. Although transcriptome and proteome can’t work as the proxy of each other [59], the question of which more accurately represents the actual functions performed by cells, the transcriptome or the proteome, should be rigorously assessed through experimental validation from multiple aspects. Metabolome quantified the metabolites’ relative level in the time points of samples collection. Although the algorithms computed the relative level of metabolites based on the key enzyme expression, the difference between the inferred and actual measurements becomes more noticeable in the non-steady state scenario of culture media. Two models’ prediction approved the succinate of SH-SY5Y is higher than other two cell lines, the inconsistency of prediction and measurement could also bring new assumption: succinate is very important for the proliferation of SH-SY5Y or the related TCA cycle genes are reprogramed in SH-SY5Y. The deeper understanding may be proposed between the different result from theoretical model and experimental observation. The essence of the phenomenon is worthy to explore further for our metabolites-transcript consistency understanding.

In an attempt to validate the potential of MGFEA on the further exploitation of public transcriptome datasets, we used MGFEA to discover novel validated metabolites from TCGA datasets and demonstrated the potential of our flexible and efficient framework. Of the four metabolites shown in figure 2c, most of them have been found to engage the progression of tumor [47,48,50,52].

For instance, we performed metabolite inference validation using the SMA dataset, which seamlessly integrates histologic data from various modalities within the MGFEA framework. Although our reference module demonstrated subtle improvement, but our attempt demonstrated constraint-based methods or flux estimation models such as scFEA [21], compass [22], METAFlux [23] and MGFEA which is compatible with high throughput single cell transcriptome datasets are also suitable for the integration of multi-omics datasets consists of MALDI, spatial RNAseq and spatial proteomics. Although the correlation between different modalities is weak, the integration of multiple modalities is also promising to produce novel knowledge and even novel research field in the future.

With the development of spatial metabolome technique [60–62], or the metabolites aptamer technique [63], it may be easier for the acquirement of the metabolic and transcript information of our interested samples, with the novel inference algorithm based on the genotype information, the understanding of interaction of genotype and phenotype could further develop and help with the human health.

In summary, MGFEA demonstrates the ability to make fast and accurate inferences about the metabolic state of a sample based on its transcriptome. It provides an algorithmic framework that can easily integrate both transcriptional and metabolic modalities from the same samples, making it a valuable tool for multimodal data integration. The further development of MGFEA can provide inspiration for the emergence of a mature integration framework across multi-omics fields, such as transcriptomics, proteomics, and metabolomics.

## Methods

### MGFEA framework

MGFEA framework (Fig. 1a) consists of data preprocess, metabolic graph integration, cell and gene embedding extraction, embedding augmentation layer and flux transformation layer.

*X*, *x_k_* : expression matrix, expression vector of kth cell

*X_r_* : matrix of spatial metabolomics matrix from spatial multimodal analysis (SMA) datasets [24]

*T^g^*, *T^c^*: the extracted Eigen vector matrix in the gene and cell dimension of the expression matrix using Principal Component Analysis

*u*, *v* : cell embedding, gene embedding

*G^m^*, *G^s^*, *A^m^*, *A^s^*: metabolic network guided gene interaction graph, spatial information graph, metabolic network guided gene interaction graph adjacent matrix, spatial information graph adjacent matrix

*φ_k_*, *θ_k_* : cell variational graph autoencoder [25] VGAE encoder parameter, decoder parameter

*φ_G_*, *θ_G_*: gene VGAE encoder parameter, decoder parameter

*F* : Flux matrix of all cells in dataset

*f* : flux vector of single cell

*S* : stoichiometry matrix of GSMM model

### GSMM model preprocess

We employed two published GSMM model, Recon3D [26] and IMM1865 [27], for MGFEA prediction of relative metabolites level. The original models have large numbers of metabolites consists of the same metabolites located in different organelles. In our modified models, we removed duplicated metabolites and used function find_blocked_reaction from python package cobrapy [28] to remove most blocked reactions.

### MGFEA metabolic interaction graph preprocess

According the metabolic stoichiometry matrix and gene co-expression relationship from expression matrix, we transformed all the information into a gene interaction graph. The graph incorporated metabolic relationships from the GSMM model, along with the intrinsic gene co-expression information in the expression matrix, as the edges between genes. For metabolic edges, we construct edges between genes connected by reaction or metabolite. For gene co-expression edges, we calculated the expression correlation of metabolic genes with highly variable genes (HVG). Edge connections are established between the top k highly variable genes and the metabolic genes with the highest correlation in their expression. To ensure the information density of HVG is comparable to that of metabolic genes, we compute the normalized dispersion of all metabolic genes. Then, the sum of normalized dispersion is used as the threshold value to select the top k HVG genes whose corresponding statistics equals to that of the metabolic genes. According to the above method, we construct a gene adjacency matrix *A^m^*.

### MGFEA embedding

MGFEA takes a preprocessed expression matrix *X* and a preprocessed GSMM model as input. Inspired by the GLUE [29] framework, we employed two separate VGAEs to learn cell embeddings in the spatial transcriptomic dataset and gene embeddings in metabolic networks separately. For the cell embedding *u*, the input consists of the expression matrix *X* and spatial coordinate information of spots *A^s^*. The obtained latent layer embedding *u* serves as the representation of different cells.

For single cell RNAseq dataset input, the training of VGAE satisfies the following loss function:

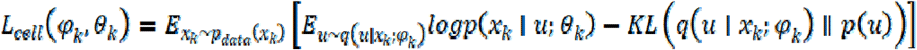

For spatial RNAseq dataset input, the training of VGAE satisfies the following loss function:

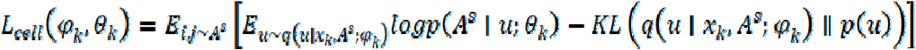

For gene embedding *v*, Using the metabolic gene interaction graph as input, the VGAE learns the intrinsic relationships of genes and acquires gene embeddings to represent different genes. The loss function of VGAE which exported gene embeddings satisfies:

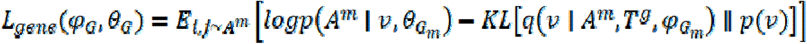

The loss of two VGAE is the latent loss of MGFEA.

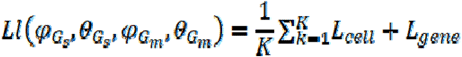

The two VGAEs learned the representations of the transcriptional cell states *u* and gene representations *v* defined jointly by the metabolic network and co-expression information. The former represents a conventional cellular state and incorporates both the expression matrix and spatial coordinate information of spots of spatial transcriptomics. The latter means the gene representations defined by gene interaction graph. The gene representations mean the genes’ location in metabolic space. The dot product of two representations shares the same matrix form as the original expression matrix. The form is used for embedding enhancement.

Taking into account the inherent projection nature of the dot product, we interpret the dot product of the two as a projection of the cellular state representation vector onto the metabolic space. For scRNAseq, by utilizing the difference between this projection and the original transcriptional expression matrix as a loss function, we enable VAE to rationalize the cell representations it learns. For spRNAseq, VGAE learns to reconstruct the spatial coordinate graph and gene interaction graph to rationalize the obtained cell embeddings and gene embeddings.

### MGFEA embedding augmentation

With the above framework, we are able to generate matrix containing both metabolic gene relationships and cell transcriptional states. Then we conducted softmax transformation between different genes within a cell based on the generated matrix. This transformation yields a matrix which contains genes’ weights in cells. Subsequently, we performed element-wise multiplication (Hadamard product) between weights matrix and the original expression matrix *X*. Through this process, we enhance the cell specific metabolic features in the original matrix *X* to preserve the transcriptional states and the augmented matrix *X_a_* improved the process of MGFEA flux-estimation.

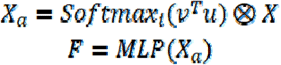

### MGFEA-Flux estimation

The balance of metabolites in reaction network is influenced by both influx and efflux. Considering that the efficiency of enzymes in metabolic networks is regulated by the regulatory genes, the transcriptional state of the cell plays a crucial role in influencing metabolic balance [30]. Building upon this premise, we utilized the transcriptional expression matrices of metabolic genes, along with a restricted set of highly variable genes, as input. We finally employed a Multilayer perceptron to estimate the fluxes of all metabolic modules (reactions).

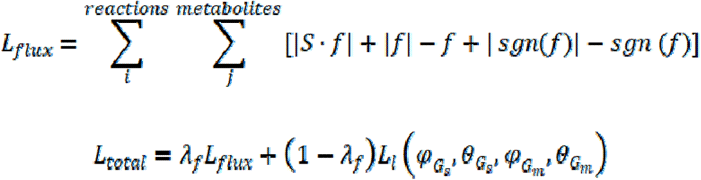

### Reference based framework

We use transcriptome and metabolome paired dataset [24] to offer a reference for spatialGraphFEA learning. We modify flux loss and add a reference loss for our model. We add reciprocal of flux to prohibit maintained decrease of flux loss. We use metabolism quantitative information as a reference and use a mean square error formula to forced predicted metabolism quantification into reference result.

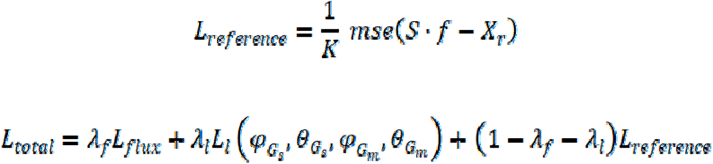

### Parameters

The weights of flux loss should be confirmed based on the epoch size. We usually used 0.5 as a default weight of flux loss. As a semi-supervised framework, MGFEA’s best parameter should be confirmed manually. When the epoch size is too big, model appears overfitting and the results even appeared as an opposite direction to the truth, we used the weight of flux loss to make flux loss converge as the training ends. We can also increase the relative weights of flux direction.

### Dataset preprocess

Expression matrix *X* is loaded in h5ad format and is normalized and log transformed. We used MAGIC [31] and STAGATE [32] for imputation. Reactions without expression in the expression matrix are removed, and the modified reaction network was used for MGFEA reaction prediction. We used highly variable genes detected by scanpy [33], spatial differential genes from spatialDE [34] and spatial correlated genes by Moran’s I from squidpy [35] for the gene interaction graph construction.

### Relative mean square error

In all instances where the relative mean square error was utilized, we first filtered out all nonoverlapping metabolites between the predicted results and the truth, we then scaled the metabolomics dataset and the metabolites’ stock level output from the model, calculating the relative mean squared error between the predicted results and the truth. For cases involving Shao metabolomics dataset, datasets from Dependency Map (DepMap) Project [36,37] and our inhouse dataset, since the vectors are not paired, so we first computed the mean metabolites level in different regions before calculating the relative mean square error for different metabolites.

### Receiver operating characteristic curve (ROC)-area under curve (AUC)

We transformed the correct direction between the different brain region pairs into the binary label. Thereby we can employed ROC-AUC metrics to assess the capability on classifying correct relative level between different brain regions of the different models. We used the true log transformed fold change between pairs of brain region as true label and the models’ predicted mean of log transformed fold change as predicted value for the ROC-AUC calculation.

### Memory usage and time consumption

We utilized python package memory-profiler(https://github.com/pythonprofilers/memory_profiler) to measure memory usage and total time consumption of different models.

### Experimental validation of Cancer Cell lines

We ordered cell lines from the vector center at Chinese institute brain research, obtaining U-87-MG (EK-Bioscience Cat.No: CC-Y1528) and HeLa (EK-Bioscience Cat.No: CC-Y1211). We acquired SH-SY5Y cell line (YC-D014) from Ubigene. We cultured cell in 90% DMEM and 10% FBS. We passaged cells every two days. The cell lines were cultured with 10cm plates. We amplified cell lines to 3-4 plates. For each cell line, once the cells reached confluence, we first removed the culture medium. We digested the cells with 0.25% trypsin for 3-5 minutes and neutralized the trypsin with 90% DMEM and 10% FBS. We pipetted to detach the cells and collected all cell mix in one 15ml centrifuge tube. After centrifuging to collect the cells, we resuspended them in PBS. Following repeated washes, we counted the cells with Countstar(Alit Biotech) and diluted them to 10^6 cells/ml. Then we separated 1ml cell suspension into a centrifuge tube and centrifuged the cells. The supernatant was removed and the pellets are stored at −80 degrees.

### RNA sequencing

The FastPure Cell/Tissue Total RNA Isolation Kit V2 (Vazyme RC112) was used to isolate total RNA from cell lines pellets. VAHTS Universal V6 RNA-seq Library Prep Kit (Vazyme NR604) was employed to generate sequencing libraries from the isolated total RNA. MGI2000 was used for sequencing the libraries. Samples are multiplexed in each lane, which yielded targeted number of paired-end, 100bp reads for each sample.

### Bulk RNA-seq data preprocess

We remove low quality reads with Trimmomatic [38], mapped reads with STAR [39] and generate counts matrix with featurecounts [40]. We used combat to remove batch effect between our in-house dataset and DepMap dataset [36,37]. The preprocessed dataset was used for subsequent flux estimation analysis.

### Metabolomics detection

We used targeted metabolomic analysis, Metabolites from the cells were extracted with 80% Acetonitrile by vigorous vortex and centrifugation at 22 000g for 20 min at 4 °C. The supernatants were used for analysis. The mix is vortexed and centrifuged. We used suspension for analysis. Chromatographic separation was performed on a I Class ultra-high-performance liquid chromatography system (Waters, Milford, Massachusetts, USA), with an InfinityLab Poroshell 120 HILIC-Z column (2.1 mm ×100 mm, 2.7 μm, agilent) and the following gradient: 0min, 100%B; 4min 84%B; 11min 40%B; 12min 40%B; 13min 100%B; 17min 10%B. Mobile phase A was 10 mM ammonium acetate in water. Mobile phase B was 10 mM ammonium acetate in 90% acetonitrile. The flow rate was 0.4 mL/min. The column temperature was kept at 35 °C and the autosampler was kept at 8 °C. The injection volume was 5 μL. Mass data acquisition of the metabolites was performed using a Triple QuadTM 7500 mass spectrometer (SCIEX, Framingham, MA) equipped with an electrospray ion source in multiple reaction monitoring (MRM) mode. The parameters of the electrospray ion source were:

neg: A: 10mM ammonium acetate, pH=8.5 B: 10% 10mM ammonium acetate, pH=8.5, 90% Acetonitrile

pos: A: 10mM ammonium formate, pH=3 B: 10% 10mM ammonium formate, pH=3, 90% Acetonitrile

The MRM transitions of all of the derivatized metabolites were shown in followed Table:

|  | metabolites | Q1 | Q3 |
| --- | --- | --- | --- |
| Group 1 | BETA-ALANINE | 90.1 | 44.2 |
| Group 2 | CITRULLINE | 176 | 159 |
| Group 3 | DEOXYADENOSINE | 252 | 136 |
| Group 4 | HYPOXANTHINE | 137 | 110 |
| Group 5 | NICOTINAMIDE | 123.1 | 80 |
| Group 6 | PROLINE | 116.1 | 70.1 |
| Group 7 | PUTRESCINE | 89 | 72 |
| Group 8 | URIDINE MONOPHOSPHATE | 325 | 97 |
| Group 9 | URACIL | 113 | 70 |
| Group 10 | FUMARATE | 115 | 71 |
| Group 11 | PYRUVATE | 87 | 43 |
| Group 12 | SUCCINATE | 117 | 73 |

### Metabolomic data analysis

SCIEX was used to process and integrate the components’ peaks and provide integrated extracted ion chromatograms for each metabolite component in all cell line samples and internal standard samples. The generated results are normalized to the internal standard samples and the normalized results are used for absolute quantification with the aid of the calibration curve.

### TCGA survival analysis

We used easyTCGA (https://github.com/ayueme/easyTCGA) to download TCGA clinical dataset and the expression matrix. We scaled the expression transcript per million (TPM) matrix using log2 transformation and utilized combat [41] to remove batch effects between glioma and GBM. After prediction of MGFEA, we divided all samples into two groups based on predicted metabolite levels and conducted Kastle–Meyer test to identify which predicted metabolite serves as a biomarker.

## Data availability

All datasets used in this study have been published and can be obtained in h5ad format from https://cellxgene.cziscience.com/datasets. The raw sequencing data of cancer cell line datasets have been deposited at CNGBdb under the accession number CNP0007635.

We used Recon3D and IMM1865 as human and mouse GSMM model. Our raw file and preprocess code can be obtained from our github site (https://github.com/Sunwenzhilab/MGFEA). Detailed message and URLs of datasets is recorded in Table 1

**Table 1.**
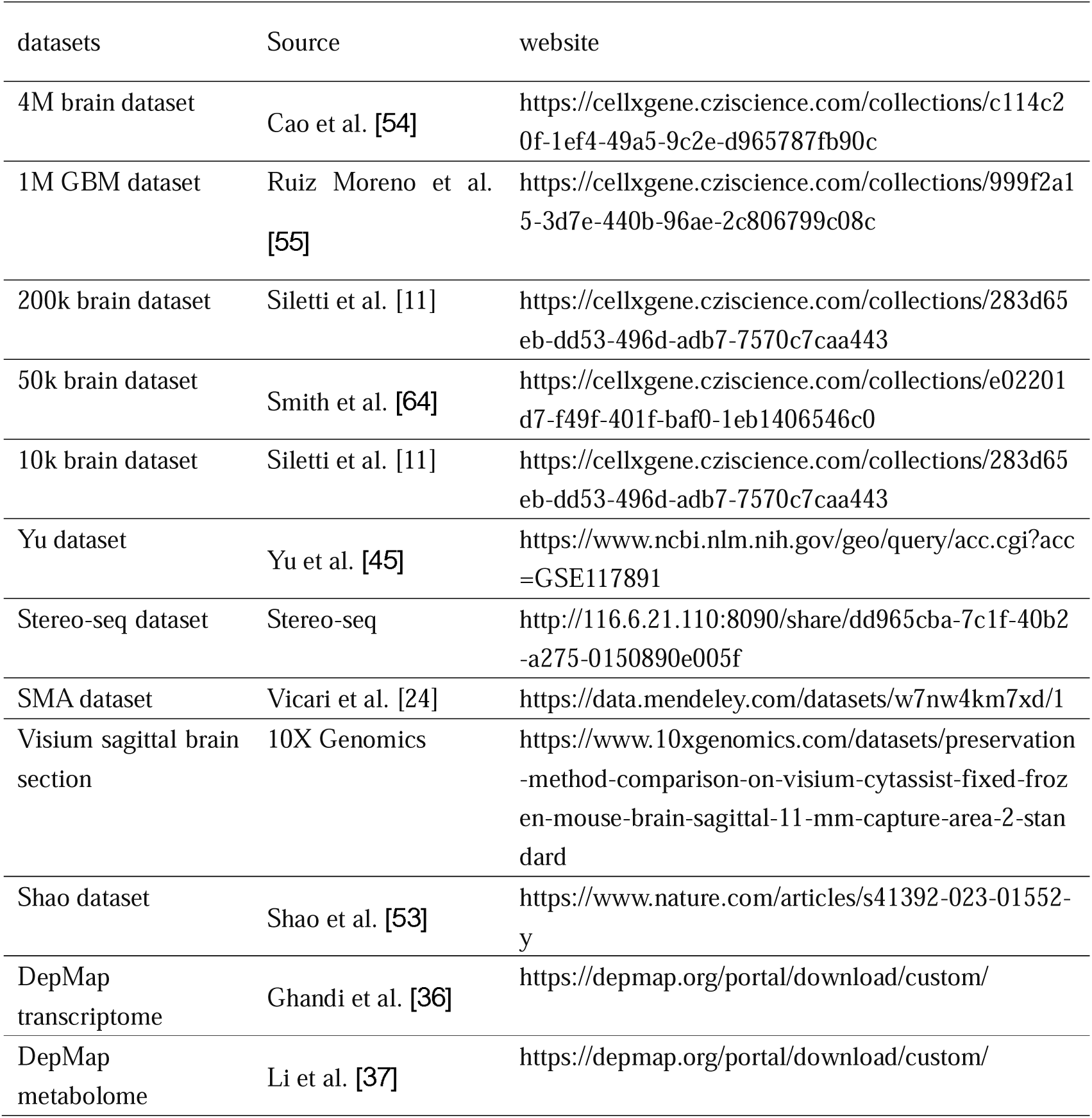

## Code availability

The code and related dataset can be accessible from the following GitHub respiratory (https://github.com/Sunwenzhilab/MGFEA).

## Author Contributions

D.A and J.L. conceived the project. W.S. supervised the whole project. J.L. implemented all model codes. J.L. and D.A. performed all data analysis. All authors reviewed and edited the manuscript.

## Acknowledgement

We thank all the member of Sun Laboratory and other colleagues from CIBR for their feedback on this work.

